# Complement in human pre-implantation embryos: attack and defense

**DOI:** 10.1101/595561

**Authors:** Martin P. Reichhardt, Karolina Lundin, A. Inkeri Lokki, Gaëlle Recher, Sanna Vuoristo, Shintaro Katayama, Juha Tapanainen, Juha Kere, Seppo Meri, Timo Tuuri

**Author notes:** These authors share senior-authorship (Seppo Meri, Timo Tuuri).

## Abstract

It is essential for early human life that immunological responses to developing embryos are tightly regulated. An imbalance in the activation and regulation of the human complement system occurs in pregnancy complications, such as pre-eclampsia and recurrent miscarriage. We hereby present the first full analysis of the expression and deposition of complement molecules in human pre-implantation embryos. Thus far, immunological imbalance has been considered in stages of pregnancy following implantation. We here show that complement activation and deposition takes place on developing human embryos already at the pre-implantation stage. Using confocal microscopy, we observed deposition of activation products such as C1q, C3 and C5 on healthy developing embryos, which highlights the need for strict complement regulation. The early embryos express the complement membrane inhibitors CD46, CD55 and CD59 and bind the soluble regulators C4bp and factor H. These findings show that complement targets human embryos, and indicate potential adverse pregnancy outcomes, if regulation of activation fails. In addition, single-cell RNA sequencing of embryos at oocyte, zygote, 4-cell and 8-cell stages showed expression of complement genes, e.g. C1s, C2, C3, C5, factor B and factor D. This shows that the embryonic cells themselves have the capacity to express C3 and C5, which may become activated and function as mediators of cellular signaling. The specific local embryonic expression of complement components, regulators, and deposition of activation products on the surface of embryos suggests that complement has immunoregulatory functions and may impact cellular homeostasis and differentiation at the earliest stage of human life.

**Statement of significance:** While canonical functions of the complement system relate to pathogen-defence, it is known to drive certain immune pathologies. The work here described shows, for the first time, expression and localization of a full range of complement molecules in human pre-implantation embryos. We demonstrate complement attack against early embryos, and show presence of embryonic defence mechanisms. Furthermore, we reveal early embryonic production of complement activators, suggesting non-canonical roles in cell signalling and development. Our findings thus reveal a fundamental role for complement at the earliest stages of human embryogenesis. Our data opens up for future studies into the role of complement, both in relation to infertility and pregnancy complications, as well as basic cellular processes during early human development.

## Introduction

The complement system is a part of the innate immune defense system, primarily involved in anti-microbial defense, clearance of debris and immune regulation. This multi-lineage enzymatic cascade functions as one of the earliest initiators of inflammation and a potent inducer of adaptive immune responses. It may be initiated through the classical and lectin pathways, driven by pattern-recognition (e.g. via C1q), or through the auto-activation of the alternative pathway. All three pathways converge at the activation of C3 to C3a and C3b, with the subsequent activation of C5 to C5a and C5b, and finally assembly of the pore-forming membrane attack complex (MAC, C5b-C9). To avoid complement attack against self-tissue, human surfaces express and/or recruit a number of membrane-bound and soluble regulators (1–3).

Increased complement activation is observed in certain autoimmune diseases, but also during pregnancy (4–6). Thus, the role of complement dysregulation has emerged as an important contributing factor to pregnancy complications and infertility, e.g. in pre-eclampsia and recurring miscarriage (7–10). Understanding how the clearance-function of complement is regulated in the context of tolerating the “semiallograft” embryo is therefore of paramount importance (11).

In addition to the manifest role of complement in immune targeting and clearance, novel functions have been revealed in recent years. Studies across multiple species have revealed unexpected roles of complement molecules in fertilization, embryonic growth and organogenesis (12). The liver is the main site for synthesis of complement components, however, these novel findings have been driven by the detection of local cell-derived complement factors and functional links to basic cellular homeostasis and metabolism (3, 9, 13–19). Furthermore, a number of animal models have revealed an effect of complement on mouse embryo hatching rate, Xenopus organogenesis as well as on rodent neuronal development (20–24).

While our understanding of local cellular complement activities is increasing, knowledge of complement expression and localization at the very early stages of human life, i.e. the pre-implantation embryonic stage, is practically non-existing. To understand the impact of complement on embryonic development, it is essential to investigate (i) potentially hazardous embryonic targeting by maternal complement (complement clearance function), as well as (ii) local embryonic production of complement components (cellular signaling and protection against maternal complement attack).

In an effort to map the localization of complement molecules and understand the role of complement activation in the early stages of human embryonic development, we here describe the local cellular expression and surface deposition of complement components using single cell RNA-transcriptomics and confocal microscopy of human non-fertilized oocytes, zygotes, 4-cell stage and 8-cell stage embryos. The current dogma in the field suggests that maternal immune tolerance towards the conceptus is induced during and after implantation. However, the findings presented here support a crucial role for innate immune mechanisms, such as complement activation, already in the pre-implantation stage of human reproduction. Furthermore, to the authors’ knowledge, these studies are the first to show the expression of multiple activating and regulatory complement components during human embryogenesis. Our findings thus suggest that complement signaling may be essential for early human development, as observed in other mammals.

## Results

### Embryos express complement regulators, activators and receptors

To understand the role of complement during early embryogenesis, we analyzed single-cell transcriptomes from embryos at different developmental stages. Included were non-fertilized oocytes, zygotes, 4-cell stage embryos and 8-cell stage embryos. Using single-cell tagged reverse-transcription sequencing (STRT), 5^’^-Transcript Far Ends (TFEs) were analyzed as described previously (25). Specifically, transcripts from the 5’-untranslated region (UTR) and upstream are reliable templates for proteins, and these were used for the analysis along with a number of reliable reads from the coding sequences (CDs). Three separate libraries were generated comparing oocyte to zygote, oocyte to 4-cell stage and 4-cell to 8-cell stage. To avoid batch-bias, these libraries were not batch corrected. We analyzed the presence of specific complement-related TFEs found in each of the four stages (Figure 1).

**Figure 1:**
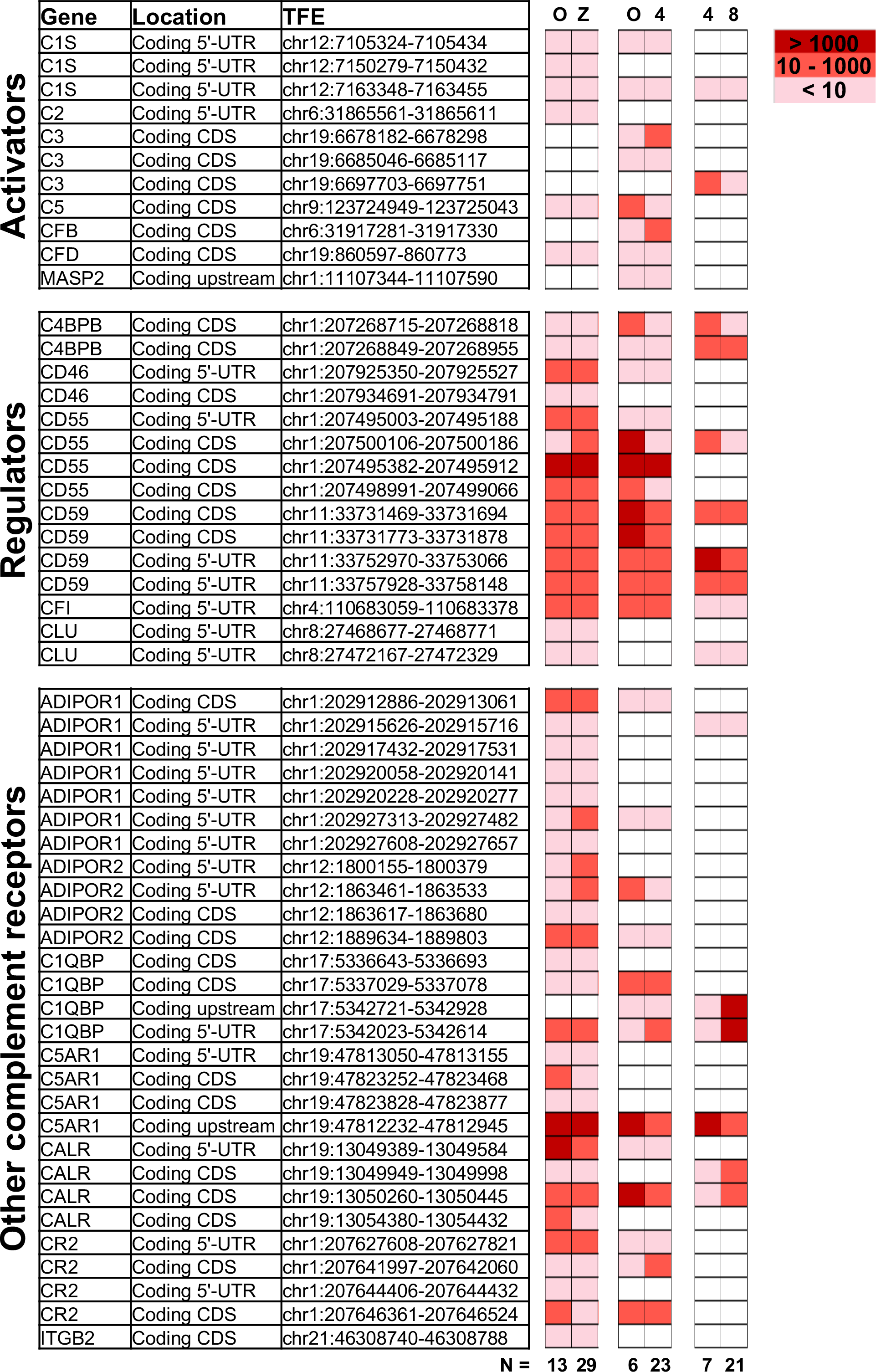
Expression of complement genes in developing embryos. Single-cell RNA sequencing was applied on oocytes, zygotes, 4-cell stage and 8-cell stage embryos to identify the expression of complement related genes during early development. Included are reads tagged to the 5’UTR or proximal region (Coding CDs), as well as known alternative splice-forms (Coding CDs). All expression levels are determined relative to spike-in reads comparing two developmental stages at a time, in three individual libraries. Expression levels are averaged over n = 6 to 29 embryonic cells, as indicated. While the data reveal the presence of non-degraded mRNA from the displayed genes, statistical analysis did not reveal significant variations in gene expression from one stage to another. For detailed functional description of identified genes see below and (1–3, 12, 26). ADIPOR1/2: adiponectin receptor 1/2, C4BBP: C4 binding protein chain B, C1QBP: gC1qR/C1q globular domain-binding protein, CALR: calreticulin receptor, CFI: complement Factor I, CLU: Clusterin, CR2: complement receptor 2, C5AR1: C5a Receptor 1, CFB: Complement factor B, CFD: Complement factor D, ITGB2: Integrin beta chain-2 (part of complement receptor 3 and 4), MASP: MBL associated serine protease.

Single-cell RNA sequencing revealed expression of a varied set of complement related genes. Transcripts correlating to a number of molecules known to protect cells from complement attack were identified. These include the membrane glycophosphoinositol (GPI)-lipid anchored complement regulators CD55 and CD59 found at all investigated stages, and CD46 not observed at the 8-cell stage. Furthermore, mRNA transcripts of soluble complement inhibitors such as C4b-binding protein (C4BP beta chain – only alternative splice-form found), factor I and clusterin were also found at constant levels from oocytes through to 8-cell stage embryo blastomeres.

In addition to mRNA transcripts of complement regulators that protect the early embryos from maternal complement attack, our data also show mRNA correlating to complement-activating molecules. The central activating components C3 and C5 are expressed by the embryos themselves already at the oocyte stage, and the transcripts remain after fertilization. In addition, proteases and cascade-components normally responsible for activation of C3, such as factor B and factor D (alternative pathway) and C1s and C2 (classical/lectin pathways) are also found. The oocytes and embryos themselves thus have transcripts for proteins that would allow cleavage and activation of both C3 and C5.

Transcripts from a number of surface receptors, commonly found to mediate activation of phagocytes and other immune cells, were also identified in the embryos. These include the C5a-receptor 1 (C5aR1), the complement receptor 2 (CR2), and integrin beta chain-2 (ITGB2/CD18 -only alternative splice-form found), which combines with CD11b or CD11c, forming the complement receptors CR3 and CR4, respectively. Transcripts of receptors for C1q (linked to clearance of apoptotic material and tissue-remodeling), such as calreticulin and C1q globular domain-binding protein (C1QBP, also known as gC1qR), were found at all investigated stages. Finally, transcripts from adiponectin receptors 1 and 2 (ADIPOR1/2), which are linked to cellular homeostasis and metabolism are also present in the early embryos.

No statistically significant increase in mRNA transcripts were identified from oocyte to the fertilized stages. The timing of active transcription is thus likely to be during oogenesis (27). Transcripts correlating to known alternative splice-forms (observed for ADIPOR1 and 2, C1QBP, C3, C4BPB, CD55, CD59, CR2, and ITGB2) may either reflect alternative protein functions, or partial breakdown of transcripts. Changes over time in the ratio between 3’UTR/degraded reads and 5’-UTR-proximal reads, may indicate an active downregulation of genes. By comparing the 3’/5’-ratios at oocyte versus 4-cell stage, we observed an increased ratio for ADIPOR2, C1QBP, C5 and CD55, suggesting their gradually increasing degradation. In contrast, a decreased ratio was observed for C3, C5aR1 and MASP2, suggesting their active transcription (data not shown).

In addition to the above-mentioned verified non-degraded transcripts, mRNA transcripts from several other complement genes were also identified in at least one of the four investigated stages. The lack of TFEs at the 5’ UTR proximal region of these reads may reflect partial breakdown of maternal transcripts, existence of promiscuous (zygotic) RNAs or novel isoforms (28, 29). Further validation is needed to determine if these transcripts produce functional protein at this developmental stage. These genes are highlighted in the Supplementary Table S1. A number of additional complement-related genes were investigated; however, no expression was detected. These are listed in Supplementary Table S2.

### Complement activation targets human embryos

Following the observation of embryonic gene expression of complement molecules, we sought to investigate if human embryos were targeted by complement activation. We considered that the complement components on the embryonic surface could originate from a combination of maternal and embryonic complement, and initially investigated deposition of the complement targeting molecules C1q, C3b, the inactivated C3d as well as C5 (Figure 2).

**Figure 2:**
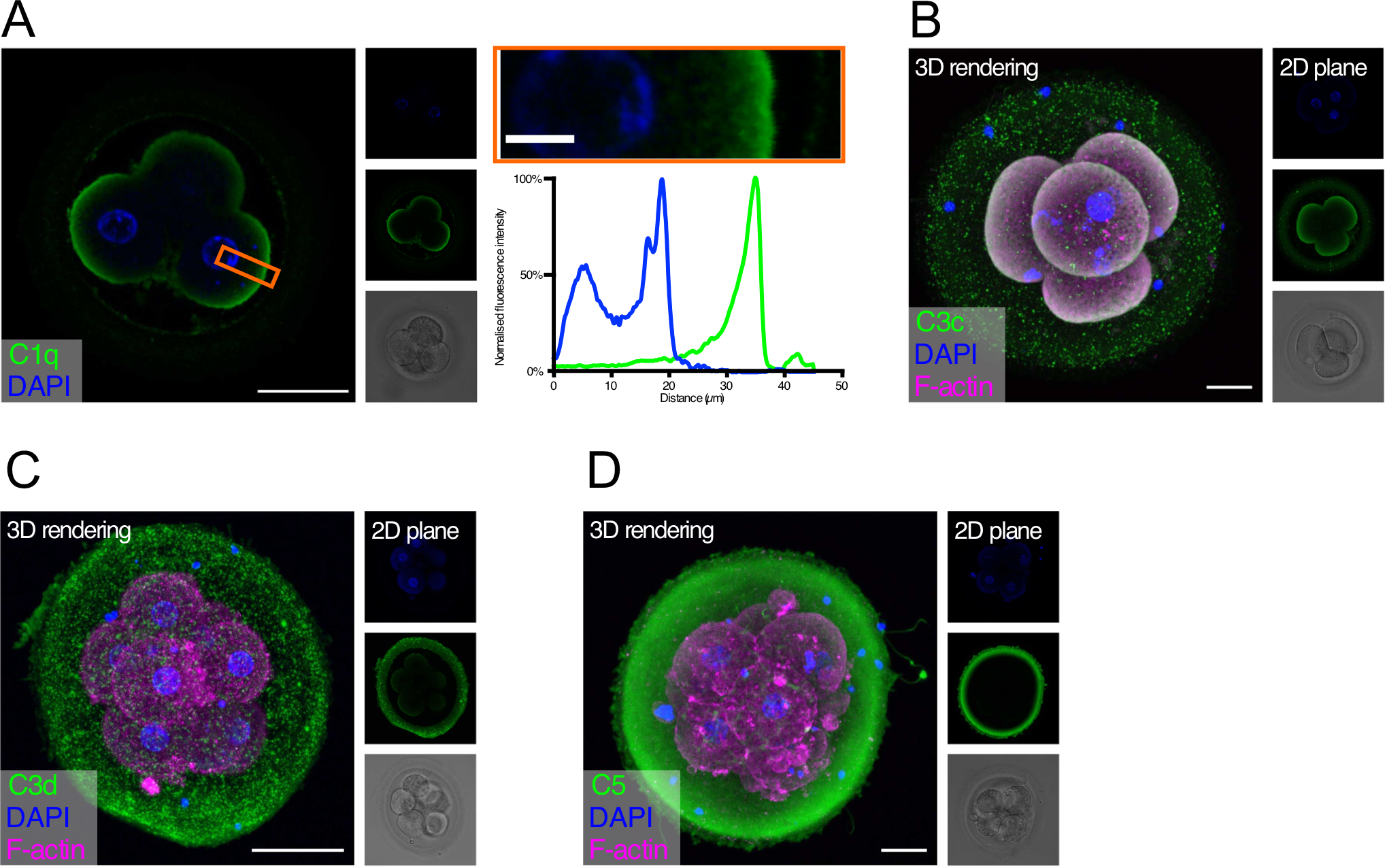
Complement targets developing embryos. Human cleavage stage *in vitro* fertilization (IVF) embryos were thawed in Vitrolife G-TL serum-free media. The embryos were then incubated with specific anti-complement antibodies and analyzed by confocal microscopy. Analysis of embryonic binding of complement activation products revealed binding of the classical pathway initiator C1q (**A**), and deposition of cascade activation components C3c/C3b/iC3b (**B**) and C3d (**C**). Finally, activation of the terminal pathway is evidenced by deposition of C5 (**D**). **A**: Left panel: Single plane, overlay of C1q (green) and DAPI (blue). Middle panels top to bottom: DAPI, C1q, and BF. Right panels: Magnification of overlay (orange insert), and below the cross-sectional distribution of fluorescence intensity. This shows C1q is specifically found on the cell surface. **B-D**: Left panels: 3D rendering, overlay of protein stain (green), DAPI (blue) and F-actin (magenta). Right panels top to bottom: DAPI, protein stain, and brightfield (BF). Scale bars: 50 μm, insert: 10 μm. For each staining, n = 3 to 4 + 1 to 3 (2PN + 3PN embryos).

Strong specific staining of C1q is visible on the cell-surface of the four-cell stage embryos (Figure 2A). No staining was observed at the cellular junctions. C1q deposition on a surface may lead to initiation of classical pathway activation. We therefore proceeded to investigate the embryonic deposition of C3 activation products. Staining for C3b/iC3b (recognized by the anti-C3c antibody, Figure 2B) shows a clear deposition on the cellular membranes. A disperse staining is also observed on the surface of the zona pellucida (ZP). The presence of C3b/iC3b on the cell membranes shows that complement activation targets human embryos. C3d is the final breakdown product of C3-inactivation and remains covalently bound to a surface long after activation has taken place. We therefore next analyzed the presence of C3d on the embryos (Figure 2C). We observe clear staining for C3d on the membranes of the embryonic cells. This shows that a large part of C3b on the embryonic surface has become degraded to C3d, indicating efficient control. In contrast to the C3b/iC3b-staining, we observed more staining for C3d on the ZP. However, variation between embryos was observed. Following deposition of complement C3b and generation of the C5 convertase, the cascade leads to the activation of C5 on the target surface. We therefore investigated the presence of C5 on the embryonic surface, using a polyclonal antibody recognizing both cleaved and non-cleaved forms of C5. Here we observed a very strong staining on the surface of the ZP, but not on the surface of the cleavage stage embryo blastomeres (Figure 2D). Despite our RNA-seq data showing cellular expression of both C3 and C5 (Figure 1), the antibodies used here did not detect any intracellular signal.

### Embryonic defense against complement attack

Complement may target ‘foreign’ as well as ‘self’ surfaces. Therefore, the presence of membrane-bound and soluble regulators is essential for preventing damage to our own tissue structures. After identifying specific complement activation on the surface of the developing embryos, and successful cleavage of C3b to C3d, we investigated the expression of the membrane-associated inhibitors CD46, CD55 and CD59 (Figure 3 and Supplementary Videos 1 and 2).

**Figure 3:**
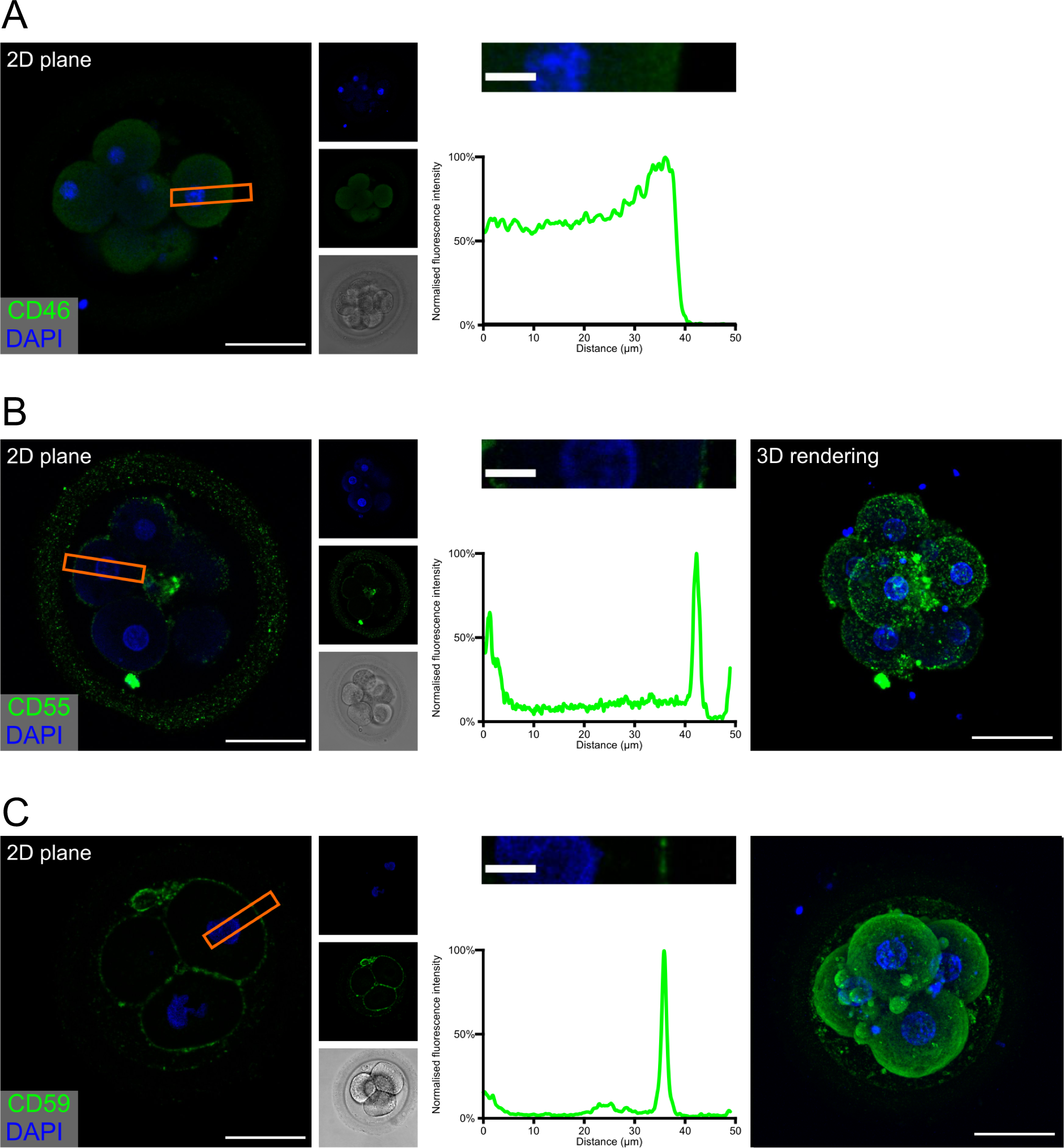
Embryonic expression of surface-tethered complement inhibitors. Human cleavage stage IVF embryos were thawed in Vitrolife G-TL serum-free media. The embryos were incubated with anti-complement antibodies and analyzed by confocal microscopy. The analysis revealed a clear staining for both CD55 and CD59, particularly at cellular junctions. In contrast, no positive signal was observed for CD46. (**A**) CD46 (**B**) CD55 (**C**) CD59. Left panels: Single plane, overlay of protein stain (green) and DAPI (blue). Second column panels top to bottom: DAPI, protein, and BF. Third column panels: Magnification of overlay (orange insert), and below the cross-sectional distribution of fluorescence intensity. Right panels (**B** and **C**): 3D rendering, overlay of protein stain and DAPI. Scale bars: 50 μm, insert: 10 μm. For each staining, n = 3 + 7 (2PN + 3PN embryos).

The expression of the three membrane regulators showed varying intensity, as expected from the gene expression data (Figure 1). In the investigated cleavage stage embryos, we observed strong staining for CD55 and CD59, but not for CD46. Interestingly, CD55 and to a lesser degree CD59, displayed a specific non-uniform localization. Both molecules are observed on the blastomere membranes. However, a stronger signal is observed specifically at the cellular junctions. While the majority of signal for CD55 is seen at the cell-cell interfaces, CD59 is found more abundantly dispersed throughout the entire cell surface. This specific pattern of CD59 expression seems to be consistent through all the early developmental stages (zygote to 8-cell stage, see supplementary Figure S1). As expected, no specific signal was observed for any of these molecules in the ZP.

In addition to the presence of membrane bound regulators, the ability to recruit soluble complement regulators is crucial for protection of viable cells against autologous complement attack (30–32). We therefore examined, whether embryos have bound the fluid phase complement regulators C4bp and factor H (Figure 4).

**Figure 4:**
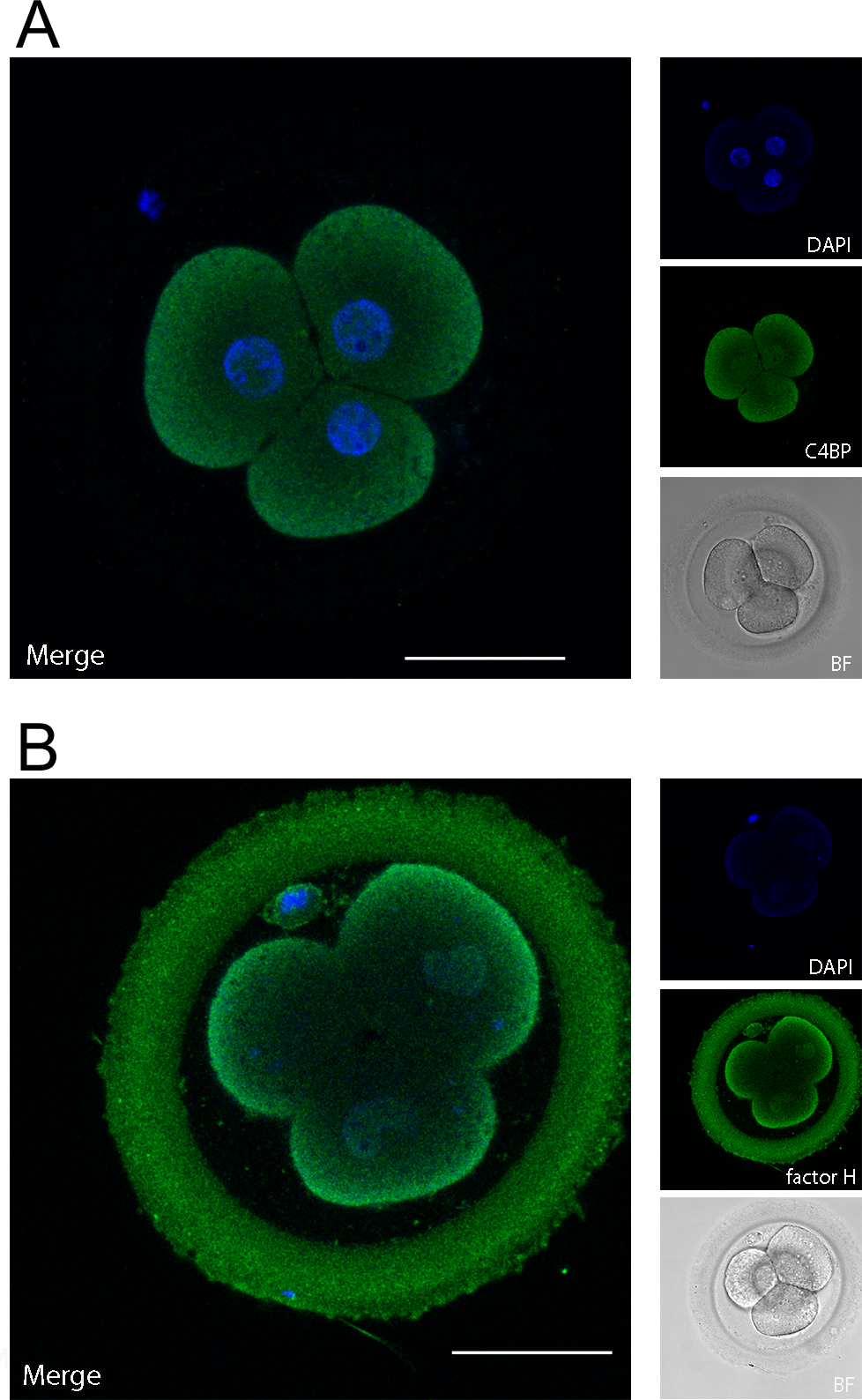
Embryonically bound soluble complement regulators. Human cleavage stage IVF embryos were thawed in Vitrolife G-TL serum-free media. The embryos were incubated with anti-complement antibodies and analyzed by confocal microscopy. Embryonic binding of the soluble complement regulators C4bp and factor H are displayed. (**A**) C4bp is recruited to the embryonic surface and show strong staining on the cell membrane. No binding is observed to the ZP. (**B**) Factor H stains both the blastomere surface as well as the ZP. Left panels: single planes, overlay of protein stain (green) and DAPI (blue). Right panels top to bottom: DAPI (blue), protein stain (green), and BF. Scale bars: 50 μm. For each staining, n = 3 + 5 (2PN + 3PN embryos).

C4bp and factor H are recruited to human surfaces immediately following complement activation, i.e. after deposition of C4b and C3b, respectively. Our data show a clear deposition of both C4bp and factor H on the cell membranes of the cleavage stage embryo blastomeres. In addition to the cellular localization of factor H, a strong staining was also observed in the ZP. This was not observed for C4bp. The binding of factor H but not of C4bp to the surface of the ZP suggests that a major part of complement activation on the ZP protein matrix (Figure 2) is driven by alternative pathway activation, which does not involve C4 cleavage. Therefore, only factor H, and not C4bp would be recruited to this surface.

## Discussion

Complement is a very potent mediator of inflammation, and untimely activation on self-surfaces contributes to a great number of pathologies, such as atypical hemolytic uremic syndrome, paroxysmal nocturnal hemoglobinuria, pregnancy disorders and kidney diseases (11, 33, 34). An understanding of the precise targeting of complement activation in various physiological settings is therefore of great importance. Animal models have been a great tool in understanding these processes. However, it is well established that variation exist between the human complement system and that of other species, such as mice (35–37). The continued investigation of human complement function is therefore crucial.

The current study is the first full analysis of the expression and localization of complement activation molecules and their regulators in human pre-implantation embryos (Figure 5). We show that complement targets the embryonic surfaces and observe deposition of complement activators, such as C1q, C3 and C5. To balance this activation, we also show expression of surface inhibitors (CD46, CD55 and CD59), as well as the deposition of soluble complement regulators, such as C4bp and factor H. This is in line with previous work showing the presence of CD55 and CD59, and possibly CD46, on human embryos (38, 39). Interestingly, the pattern of CD55 and CD59 expression at cellular junctions suggests that these molecules, in addition to complement regulation, may be directly involved with cellular interactions, such as signaling or adherence. While being GPI-anchored to cell membrane rafts or caveolae CD55 and CD59 would be in a position to transmit robust activating signals to cells.

**Figure 5:**
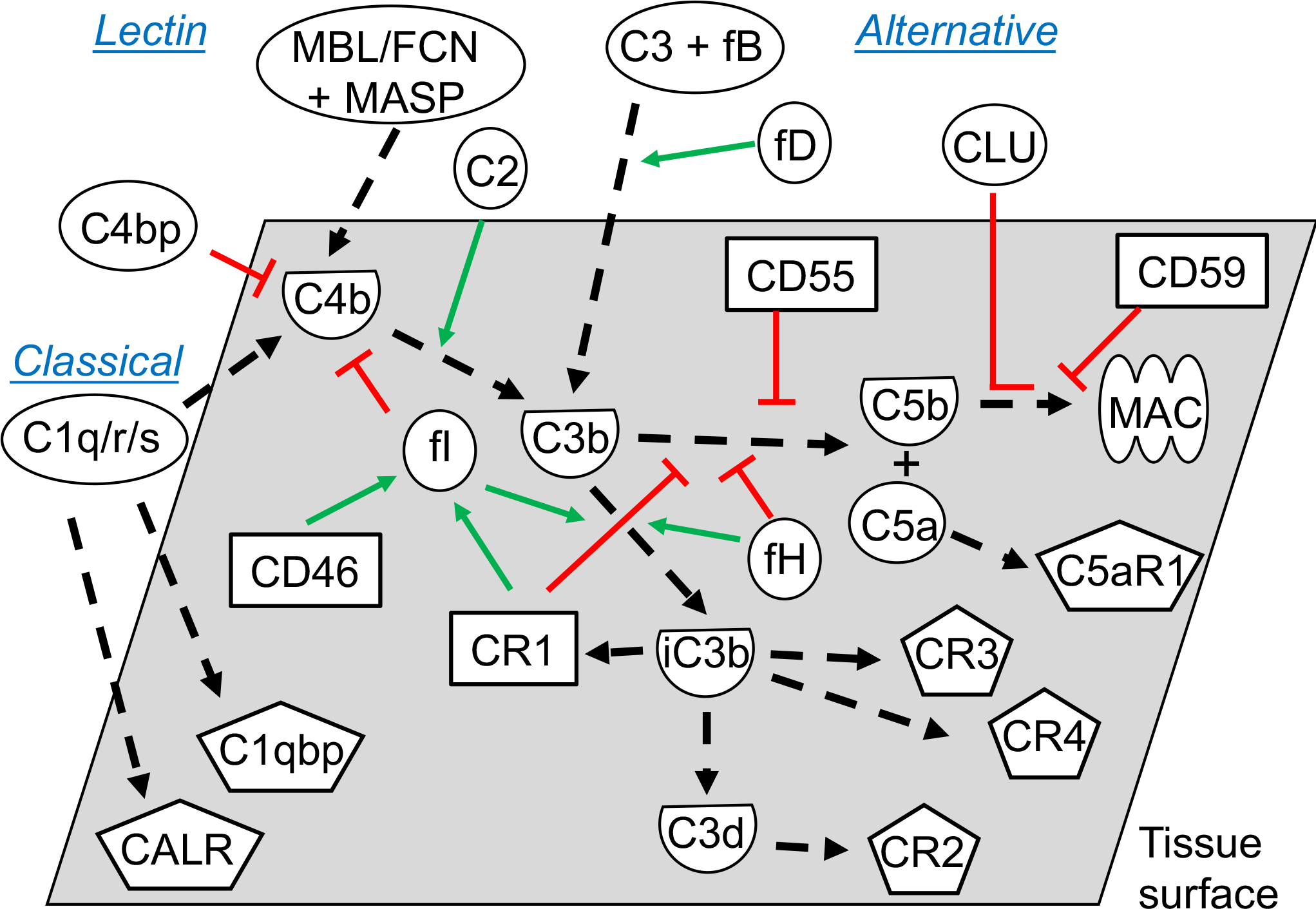
Functional overview of the embryonic complement system. Indicated are the canonical functional roles of membrane-expressed complement regulators (squares), soluble complement components (circles), and their cleaved activated membrane-deposited forms (demi-circles). Finally, embryonically expressed complement receptors are shown (pentagons). All molecules depicted were found to be expressed or bound by the embryos in this study (exceptions: C4, mannose binding lectin (MBL), ficolins (FCNs), and some MAC-components). The classical and the lectin pathways are initiated by target-binding of pattern recognition molecules such as C1q, or MBL and FCNs, respectively. Utilizing their associated proteases C1r/s or MASPs, they activate C4 and C2, which subsequently activate C3. Alternative pathway activation of C3 occurs when factor D cleaves C3-associated factor B, which generates a novel C3-cleaving enzyme; C3bBb. Cleavage of C3 by either pathway leads to generation of soluble C3a and surface-deposited C3b, which amplifies alternative pathway C3 activation, and subsequently activates C5 to C5a and C5b. Finally, C5b initiates the assembly of the pore-forming MAC (1, 2, 40). To avoid excessive immunological targeting of self, human cells express or recruit inhibitors of complement activation. Membrane regulators may function; 1) by disrupting the enzymes cleaving C3 and C5 (CR1, CD55), 2) as co-factors for factor I-mediated degradation of C3b and C4b (CR1, CD46), or 3) by inhibiting MAC-formation directly (CD59). Soluble regulators such as factor H and C4bp inhibit activation by mechanisms 1 and 2, while clusterin work through mechanism 3. Inactivation of C3b, leads to generation of iC3b, C3dg and finally C3d. While C3b and iC3b function as opsonins for increased phagocytosis by antigen presenting cells (through CR3 and CR4), C3d has important biological functions as an important internal adjuvant aiding antigen uptake by dendritic cells and inducing efficient antibody responses in B cells (through CR2). (41–44). While CR2, CR3 and CR4 expression is mainly described on immune cells, our study found embryonic expression of these receptors, along with the signaling receptors for C1q; calreticulin (CALR) and C1qbp.

Single-cell RNA sequencing revealed that mRNA from a number of complement components are found at various stages of early development, i.e., in oocytes, zygotes, as well as at the 4-cell stage and 8-cell stage blastomeres. Our analysis identified validated 5’ UTR-proximal TFEs (a reliable indicator for protein translation), or alternative splice-forms of transcripts from the genes ADIPOR1, ADIPOR2, C1QBP, C1S, C2, C3, C4BPB, C5, C5AR1, CALR, CD46, CD55, CD59, CFB, CFD, CFI, CR2, ITGB2 and MASP2. The CD-transcripts representing alternative splice-forms may have unknown functional relevance at the embryonic stage. The role of these TFEs require further validation. Comparing oocyte to the 4-cell stage, we observed increased degradation of ADIPOR2, C1QBP, C5 and CD55, and decreased degradation of C3, C5aR1 and MASP2 (measured by the 3’/5’ ratio). Though no statistically significant increase in transcription was detected, a decreased degradation supports a role in helping the embryo on its way to the uterus. In contrast, increased degradation suggests a primary function at the oocyte stage. In addition to the 5’UTR-proximal reads, we identified partially degraded transcripts or potentially novel isoforms from additional complement genes. These include genes encoding important molecules such as ADIPOQ, C1q-C, C1R, C7, CDH13, CR1, and CFP. While our method cannot distinguish between explicit embryonic expression or earlier maternal expression of these partially degraded mRNAs, the presence of these gene-transcripts suggest expression during oogenesis. Proteins translated during oogenesis may be important at this stage only, but are also likely to remain in the early fertilized embryo. Combined with the validated 5’-UTR and splice-form reads, we thus demonstrate that human oocytes and pre-implantation stage embryos produce a very wide range of complement molecules.

### Immunological targeting of human embryos

By confocal microscopy we observed targeting of human embryos by complement. While the potent inflammatory role of complement has mainly been studied in the context of serum, it is well established that complement components are found in mucosal secretions, e.g., from cervix, uterus, and fallopian tubes (22, 45–49). The presence of complement in the uterine compartment, alongside the data presented here, indicates that human embryos are targeted by complement in a physiological setting. This makes the observed expression of surface inhibitors and the recruitment of soluble regulators essential for the survival of the embryo already prior to implantation. Numerous links between complement and pregnancy complications have been described in the literature, all related to implantation, placentation or later development (4–7, 9–11). Our data provide a novel, much earlier, mechanism, whereby a faulty or insufficient complement regulation may predispose to pregnancy disorders and miscarriage.

The expression of CD55 and CD59 throughout the investigated period of embryogenesis, and CD46 at certain stages, may be important for embryo survival. However, attempts of stem-cell transplantation show the importance of soluble regulators for cell survival as well (30, 31). Our data reveal presence of clusterin and factor I mRNA, and deposition of C4bp and factor H on the cleavage stage embryos. The inhibitory effects of these molecules were substantiated by our staining for C3-degradation products. Importantly, only the non-degraded C3b will lead to continuation of the complement cascade and formation of the C5 activating convertase. No staining was observed for C5 on the blastomere membranes, thus showing that the kinetics of the complement regulation favor degradation of C3b deposited on the cell surfaces. However, on the ZP, the lack of membrane regulators may favor a different outcome. The presence of factor H, but not of C4bp, on the surface of the ZP suggests that the majority of complement activation against the ZP protein matrix is a result of alternative pathway activation, which does not involve C4 cleavage. A strong C5 deposition was observed on the ZP, indicating that this layer absorbs the most intense complement attack. Though our staining for C3d revealed that a lot of the C3b deposited has become degraded, the kinetics of C3b-degradation versus C5-convertase formation has still favored C5 activation. Given the stronger deposition of complement activation products, a function of the ZP may be to divert complement from the cell membranes, and act as a protective layer for the developing embryo also in this respect. The role of factor H as the main inhibitor of complement activation against the ZP, highlights the potentially detrimental impact of factor H deficiencies on reproduction. Still, while we here observe potentially harmful pro-inflammatory stimuli in response the pre-implantation embryo, the impact on pregnancy outcome may be less clear. Local inflammatory processes are crucial for decidualization of the endometrium following implantation and in establishing immune tolerance towards the developing conceptus (50). While immunological opsonization and clearance of the conceptus would be catastrophic, the activation of complement may therefore also contribute to priming the immunological landscape following implantation.

### Cellular complement activation and signaling

From the perspective of the developing embryo, the expression of complement regulators and binding of inhibitors is useful for protection against the clearance function of complement. However, our data show that complement activation components are also expressed. This suggests other functional roles of complement, e.g. in cellular signaling or in metabolism as has been suggested for a wide range of cells, including stem cells (12, 51, 52). With the local expression of C1s, C2, C3, C5, factor B and factor D, the oocyte and potentially later embryonic cells themselves produce the molecules necessary for initiating complement. Activation of cell-derived C5 and subsequent autocrine binding to its receptors C5aR1 and C5aR2, has been shown to initiate a number of cellular signaling events (51, 52). As we also identify expression of C5aR1, it is possible that complement is utilized for cellular signaling events, as has been suggested for human stem cells (53–55). It has previously been shown in the human oviductal epithelium, that the combined expression of molecules such as factor B and factor I, together with C3 is enough to produce an active C3-convertase and generate C3-cleavage products such as iC3b (22). The study by Tse et al. revealed an embryotrophic effect of iC3b on mouse embryos. Our data show that expression and/or deposition of the molecular machinery mediating the embryotrophic effects in mice, also exists in humans.

### C1q and tissue-remodeling

While the gene expression data suggest that C1q and C1r may be expressed in oocytes, they appear to be degraded later. However, the staining data show a clear binding of C1q to the blastomere membranes. The strongly expressed C1qbp may function as an essential regulator of complement activation at this stage (56). Mouse studies have indicated C1q as an important signaling molecule in stem cell differentiation, and later during development as a crucial molecule for tissue organization (24, 57, 58). Furthermore, human studies have related altered C1q-mediated clearance of debris and apoptotic material from the placenta in pre-eclampsia (9). Calreticulin, of which we found mRNA transcripts at all investigated stages, is thought to act as a receptor for C1q and collectins to mediate clearance of apoptotic materials. The deposition of C1q observed in the current study and the high levels of mRNA from complement-modulatory proteins in the early embryo suggest that C1q may have similar functions already at this developmental stage.

In conclusion, we here provide evidence for complement targeting of early human embryos, along with a substantial expression and/or recruitment of complement inhibitors. These findings suggest that a lack of appropriate inhibition of the activation cascades, and the generation of C3 and C5 cleavage products on the pre-implantation embryonic tissue may be detrimental to the developing embryo and the inflammatory state of the maternal endometrium. However, the functional relevance of complement-deposition on pre-implantation embryos is very likely to extend beyond mis-directed immunological clearance. Our finding of early cellular expression of complement-activating molecules supports emerging roles of complement in basic cellular processes, such as metabolism and differentiation. This is the first study to identify the extent of complement involvement in the pre-implantation developmental stage in a bona fide human model. The data presented here thus highlight the importance of further studies into the role of complement, both in relation to fertility and pregnancy complications, as well as in relation to basic cellular processes during early human development.

## Materials and Methods

### Ethical considerations

Oocytes and embryos utilized for the single-cell RNA sequencing were collected in Switzerland and Sweden. All analyses were performed in Sweden. The full protocol was approved by the ethical committees in Switzerland (authorization CE2161 of the Ticino ethical committee, Switzerland) and in Sweden (Dnr 2010/937–31/4 of the Regional Ethics Board in Stockholm). Embryos used for confocal imaging were donated by patients at the Helsinki Women’s clinic Fertility unit, Finland, after informed consent. The study was approved by the local ethics committee (124/13/03/03/2015 and DNr 308/13/03/03/2015). All cells and embryos were generated for the sole purpose of IVF. Following standard procedures embryos generated for IVF treatment, but not immediately transferred to the uterus, were cryopreserved. Upon termination of the freezing contract, embryos were either discarded or donated for research purposes. Abnormally fertilized triploid (3PN) embryos were donated at fresh cycles and used either fresh or after vitrification and warming at later time point.

### Reanalysis of single-cell RNA sequencing data on human oocytes and blastomeres in the preimplantation embryos

Initial transcriptomics data were generated earlier (25). Embryos utilized for single cell-RNA sequencing were collected and cultured as described. Individual blastomeres were obtained by laser-assisted biopsy, the ZP was removed and cells were placed in lysis buffer in individual wells of a 96-well plate. For single-cell RNA sequencing STRT was applied (59). Three individual libraries were prepared from the total number of single cells; oocytes: 19 cells, zygotes: 29 cells, 4-cell stage blastomeres: 30 cells and 8-cell stage blastomeres: 21 cells, as described (25). Expression levels were correlated to eight synthetic spike-in RNAs (ArrayControl RNA spikes Ambion, cat. no. AM1780) (60). Following amplification, the synthesized cDNA was sequenced on the Illumina platform, filtered, demultiplexed and trimmed as described (25). Estimation of the ratio of transcripts per cell was done by comparing total reads to total spike-in RNA associated reads (all with sample-specific barcodes). Following pre-processing, the reads were aligned to human UCSC genome hg19, ArrayControl RNA spikes and human ribosomal DNA complete repeat unit (GenBank: U13369) by TopHat version 2.0.6 (61) and annotated by genomic features. The aligned STRT reads were assembled by sample types using Cufflinks (61) and counted as TFEs, as described (25).

### Complement gene-expression analysis

A list of genes relevant to complement function was generated based on an exhaustive analysis of functional studies in the field (1–3, 12, 26). The method of TFE-based quantification is implemented as open-source software (https://github.com/shka/STRTprep). This method was used to identify expressed complement genes. In brief, the TFEs were defined by STRT RNAseq reads, which correspond to the 5’-end of polyA-tailed RNAs. Therefore, TFEs not at the 5’-UTR or the proximal upstream were excluded from the investigation of known protein-coding genes, as mRNAs suggested by these TFEs were less likely to have a methionine for translation of functional protein. Furthermore, TFE-based quantitation provides an advantage over normal sequencing approaches particularly in studies of preimplantation development, where the gene-based quantitation methods also sum promiscuous (zygotic) RNAs and degraded (maternal) RNAs, which are less likely to translate into proteins (28, 29). Therefore, the tagging of RNAseq reads to the 5’UTR was a positive selection criterion for expression. The integrity of the mRNA reads not directly tagged to the 5’-UTR in human GRCh37/hg19 build was assessed by the Zenbu Genome browser, the UCSC Genome Browser and the GENSCAN online tools (62, 63). This led to the inclusion of a number of reads tagged to the CDs of coding transcripts. Specifically, sequences tagged a few codons downstream of the 5’ end were included. Also, sequences corresponding to alternative splicing events and sequences that aligned with expressed spliced human sequence tag reads within the target gene were included in the analysis.

### Statistical analysis for differential expression

The R package pvclust (64) was applied to exclude outlier samples. Subsequently, differential expression levels were tested by SAMstrt (65), a version of SAMseq (66) modified for spike-in-based normalization.

### Collection and culturing of cleavage-stage human embryos for confocal imaging

Cleavage stage embryos utilized for confocal microscopy were cultured to 4- or 8-cell stage embryos in a sequential culture system (G-IVF/G-1PLUS, Vitrolife) at 37°C and 5% CO2, 5% O_2_. Embryos were frozen and thawed using Vitrolife FreezeKit Cleave and ThawKit Cleave, respectively (Vitrolife Sweden AB, SE-421 32 Västra Frölunda, Sweden). After thawing, embryos were cultured in Vitrolife G-TL medium until processed for immunostaining.

### Confocal microscopy

Embryos were fixed with 4% paraformaldehyde in phosphate-buffered saline (PBS) solution for 15 min at room temperature, washed with PBS and permeabilized using 0.5% Triton^®^ X-100 (Fisher Scientific, Geel, Belgium) in PBS for 15 min, and blocked for unspecific staining using Ultra Vision Protein Block (Thermo Scientific, MI) for 8-10 min, all at room temperature. Primary antibodies, mouse anti-CD46 (GeneTex Inc., Irvine, California, US; 1:100), mouse anti-CD55 (IBGRL Research Products, Bristol, UK; 1:100), mouse anti-CD59 (IBGRL Research Products, Bristol, UK; 1:100), goat anti-factor H (Calbiochem, La Jolla, California; 1:100), mouse anti-C4bp (Quidel, San Diego, California; 1:100), rabbit anti-C1q (DAKO Denmark A/S, Glostrup, Denmark; 1:100), goat anti-C5 (Cappel, Organon Teknika Corp., West Chester, Pennsylvania; 1:300), rabbit anti-C3d (DAKO Denmark A/S, Glostrup, Denmark; 1:100), rabbit anti-C3c (DAKO Denmark A/S, Glostrup, Denmark; 1:100) were diluted in PBS + 0.1% Tween20 (Fisher Scientific, Geel, Belgium) and incubated over night at +4°C. After three washes in PBS + 0.1% Tween20, embryos were incubated 2h at room temperature while rocking in secondary antibody donkey anti-mouse AlexaFluor^®^594, donkey anti-goat AlexaFluor^®^488, or donkey anti-rabbit AlexaFluor^®^488 (all from Thermo Fisher Scientific; 1:500) diluted in PBS + 0.1% Tween20. F-actin was stained with AlexaFluor^®^647 Phalloidin (Thermo Fisher Scientific; 1:100) and nuclei were stained with DAPI (Thermo Fisher Scientific; 1:500). Images of the embryos were acquired using an inverted TCS SP8 MP CARS confocal microscope (Leica Microsystems, Mannheim, Germany) and Leica HC PL APO CS2 40x/1.10NA water and Leica HC PL APO CS2 63x/1.20NA water objectives.

### Confocal image processing

Confocal images were processed using Fiji (http://fiji.sc) and Imaris (BitPlane, Oxford Instruments). Depending on the dataset, preprocessing consisted of applying a sliding-window averaging in the z dimension, denoising with Rolling Ball and smoothing by applying either a Gaussian filter or a 3D-Median filter (kernel 1). When necessary, DAPI signal was isolated from the background by applying a binary mask (obtained by local maxima functions) to the raw image to specifically render nuclei only. Fluorescence intensity profiles were obtained by averaging lines cropped in the inset (to smooth the noise) and are plotted as normalized to the maximum peak. 3D renderings were obtained using Imaris.

## Acknowledgements

We are grateful to the anonymous donors of cells enabling this study. The computations were performed on resources provided by SNIC through Uppsala Multidisciplinary Center for Advanced Computational Science (UPPMAX) under Project b2010037. Confocal microscopy was carried out at the Bioimaging Unit, University of Helsinki, Finland. Funding for the project was provided by the Finnish Cultural Foundation, the Jenny and Antti Wihuri foundation, the Academy of Finland, Helsinki University Hospital Funds, Foundation ARC for cancer research (grant #20171206504), Knut and Alice Wallenberg Foundation, Swedish Research Council, Sigrid Jusélius Foundation and Jane and Aatos Erkko Foundation. GR is a member of the CNRS ImaBio GdR.

## Author contributions

M.P.R., J.T., J.K, S.M. and T.T. designed research; M.P.R., K.L. and S.K. performed research; M.P.R., K.L., A.I.K, G.R. and S.V analyzed data; and M.P.R., K.L., A.I.K. and S.M. wrote the paper.

## Abbreviations

CDs: coding sequences
IVF: *in vitro fertilization*
STRT: single-cell tagged reverse-transcription
TFE: transcript far end
ZP: zona pellucida
5’-UTR: 5’-untranslated region

## Complement in human pre-implantation embryos: attack and defense

**Supplementary Table 1:** Complement gene-transcripts mapped to coding regions outside the 5’ UTR. Highlighted here are additional complement-related transcripts mapped to coding regions outside the 5’ UTR. Based on the time-dependent regulation of gene transcription surrounding the human genome activation, it is not possible to determine the specific expression of these genes at the various stages. The massive presence of certain genes through all tested stages does however indicate that gene transcription from particular loci is active during the investigated early embryonic stage. C1q-C: C1q, C-chain, CR1: complement receptor 1.

| Partially degraded transcripts / potential novel isoforms |  |  |  |
| --- | --- | --- | --- |
| Oocyte | Zygote | 4-cell | 8-cell |
| Adiponectin | Adiponectin | Adiponectin | Adiponectin |
| C1q-C | C1q-C | C1q-C | C1q-C |
| C1R | C1R | C1R | C5 |
| C7 | C3 | C7 | C7 |
| Cadherin13 | C7 | Cadherin13 | CD46 |
| CR1 | Cadherin13 | CD46 | CR1 |
| Properdin | CR1 | CR1 | Factor D |
|  | Factor B | Properdin | Properdin |
|  | Properdin |  |  |

**Supplementary Table 2:**
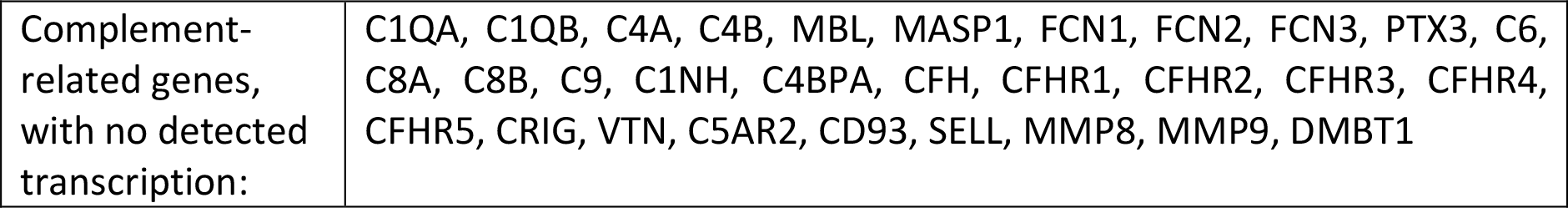
Complement genes with no expression. Listed are complement related genes not found to be expressed. MBL: Mannose binding lectin, MASP: MBL associated serine protease. FCN: Ficolin, PTX: Pentraxin, C1INH: C1 inhibitor, CFH: Complement factor H, CFHR: Complement factor H related, CRIG: Complement Receptor of the Immunoglobulin superfamily, VTN: Vitronectin, VSIG4, C5AR2: C5a Receptor 2, SELL: L-selectin MMP: matrix metalloproteinase, DMBT1: deleted in malignant brain tumor 1 (also known as SALSA or gp340).

**Figure S1:**
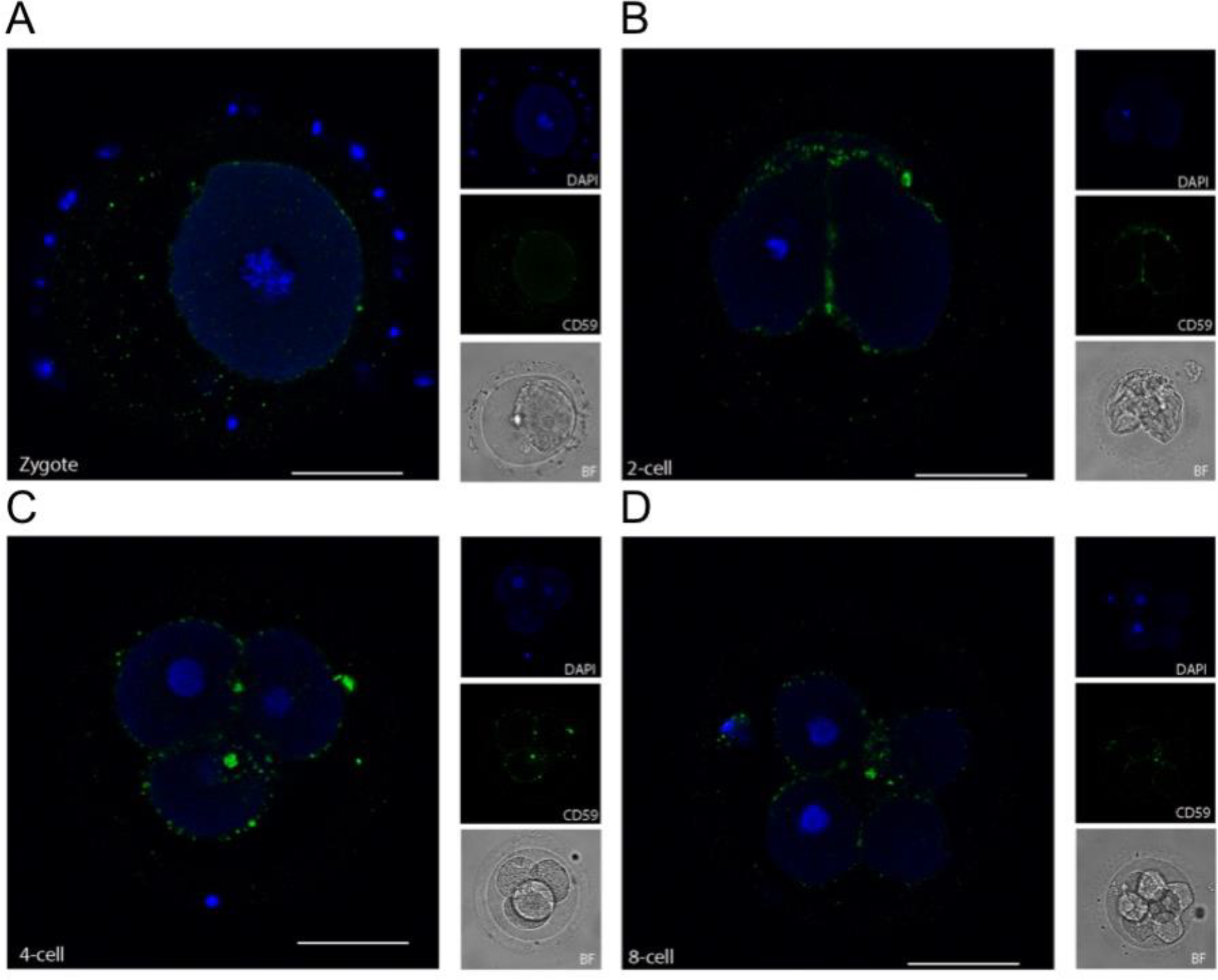
Localization of CD59 from zygote through to 8-cell stage. To support the observation of CD59 expression at all developmental stages, we stained embryos at zygote through to 8-cell stage for this marker. At the zygote stage (**A**) CD59 stains evenly on the membrane surface. As the embryo develops through 2-cell stage (**B**), 4-cell stage (**C**), and 8-cell stage (**D**), the staining gather into particular clusters concentrated at the cellular junctions (**B**-**D**). Displayed are 2PN embryos (3 PN for the zygote). Left panels: single planes, overlay of protein stain (green) and DAPI (blue). Right panels top to bottom: DAPI (blue), protein stain (green), and BF. Scale bars: 50 μm. Due to the very limited material, these stainings were only performed with n = 1 2PN embryos per stage. Zygote, plus additional 4 3PN embryos.

### Supplementary videos

**Video 1:** Human cleavage stage IVF embryos were thawed in Vitrolife G-TL serum-free media. The embryos were incubated with anti-complement antibodies and analyzed by confocal microscopy. The analysis revealed a clear staining for CD55, particularly at cellular junctions. 3D rendering, overlay of CD55 (green) and DAPI (blue). Scale bar as indicated.

**Video 2:** Human cleavage stage IVF embryos were thawed in Vitrolife G-TL serum-free media. The embryos were incubated with anti-complement antibodies and analyzed by confocal microscopy. The analysis revealed a clear surface staining for CD59, with increased signal at cellular junctions. 3D rendering, overlay of CD59 (green) and DAPI (blue). Scale bar as indicated.

